# Acyl plastoquinol is a major substance that co-migrates with triacylglycerol in cyanobacteria

**DOI:** 10.1101/2022.11.08.515603

**Authors:** Natsumi Mori-Moriyama, Toru Yoshitomi, Naoki Sato

## Abstract

Various studies have suggested the presence of triacylglycerol in cyanobacteria, but no convincing evidence exists. We purified a substance co-migrating with triacylglycerol and determined its structure using mass spectrometry, gas chromatography, and ^1^H- and ^13^C-NMR. The major components were palmitoyl and stearoyl plastoquinols (acyl plastoquinol). Acyl plastoquinol has never been described before. The level of acyl plastoquinol was 0.4% of the total lipids. We still do not have clear evidence for the presence of triacylglycerol. If present, the maximum triacylglycerol level, must be at most 10% of acyl plastoquinol or less. The *Synechocystis* Slr2103 protein was suggested to synthesize triacylglycerol, but the product could be acyl plastoquinol. The possible roles of this novel compound in photosynthesis should be a new focus of research.

## Introduction

Triacylglycerol (TAG) is a common storage lipid in eukaryotes. In the current trend of green energy development, exploiting photosynthetic energy in oil production is a good goal of biotechnological research. Accordingly, the algal production of TAG has been a focus of technological advancement for these decades. We were also involved in the studies on TAG accumulation in *Chlamydomonas reinhardtii* and *Cyanidioschyzon merolae* (Moriyama et al. 2018, Toyoshima et al. 2016). Cyanobacteria have also been studied in the framework of algal oil production. However, it is still challenging to achieve a high-level accumulation of TAG in cyanobacteria (see, e.g., Tanaka et al. 2020). We must understand that cyanobacteria are prokaryotes and distinguish them from eukaryotic algae. Some bacteria accumulate TAG to several tens of percent of dry weight (for a review, see Alvarez and Steinbüchel 2002). In algae, TAG accumulates in the lipid droplets in the cytosol in close contact with either the endoplasmic reticulum or chloroplasts. TAG is also present in the plastoglobules as a minor component, but physically distinct lipid droplets are present only outside the chloroplasts (Moriyama et al. 2018).

TAG has repeatedly been reported from cyanobacteria for decades. Old reports described TAG as a constituent of cyanobacteria in natural habitats (Taranto et al. 1997, Řezanka et al. 2012). Paramuna and Sumners (2014) gave a more rigorous analysis of the pure culture of cyanobacteria. They detected, by fluorescence microscopy, minute lipid droplets in the cytoplasm of *Nostoc punctiforme*. They also found a lipid spot co-migrating with authentic TAG in TLC analysis. The materials contained mainly saturated fatty acids in the exponential growth phase, but unsaturated fatty acids dominated the stationary phase. The authors related the material to the lipid droplets detected by microscopy.

The endosymbiotic hypothesis of chloroplast origin is often used to explain some aspects of the similarity of chloroplasts and cyanobacteria. The fibrillin (also called plastoglobulins) homologs in cyanobacteria were taken as evidence for the presence of a cyanobacterial counterpart of plastoglobule (Lohscheider and Bártulos 2016). The acyltransferase PES1 synthesizes phytyl esters in the plastoglobules (Lippold et al. 2012). A cyanobacterial acyltransferase Slr2103, a homolog of PES1, was reported to synthesize TAG rather than phytyl esters (Aizouq et al. 2020). This study was believed to open a new study of cyanobacterial TAG (Santana-Sánchez et al. 2021). However, many researchers have already noticed that the data in Aizouq et al. (2020) were strange. Notably, the TAG level was too high (comparable to the chlorophyll level) if the description was understood literally. However, the TLC spot of putative TAG was smaller than the free fatty acid spot.

In this respect, the current status of cyanobacterial TAG is confused. Some works claim to detect TAG in cyanobacteria, whereas serious doubt about the reality of cyanobacterial TAG remains as exposed by e.g., Tanaka et al. (2020). Doubts have been expressed as oral or poster presentations in some academic meetings (e.g., Mori and Sato, poster PO-59 in the International Symposium of Plant Lipids. Yokohama, Japan, 2018), but have rarely been presented as written papers except Tanaka et al. (2020).

In the present study, we determined the structure of the major component of the spot that co-migrates with TAG in cyanobacteria. We identified acyl plastoquinol (APQ), a novel compound that has never been described in biological literature. We also tried identifying TAG in cyanobacteria, but the maximum TAG level must be one order less than the level of APQ. The enzymatic reaction catalyzed by the product of the *Synechocystis* slr2103 gene could produce APQ rather than TAG. APQ could function in regulating photosynthesis rather than a storage material like TAG. We will discuss the possible roles of APQ in cyanobacteria.

## Results

### Efforts to detect TAG

We developed a convenient but reliable method of detecting a minute amount of TAG in cyanobacteria. Previously, Aizouq et al. (2020) used a solid-phase extraction system to enrich non-polar lipids from the total extract of cyanobacteria. However, we preferred to use conventional two-dimensional thin-layer chromatography (TLC) to separate TAG while preserving all other lipid components on a single plate (**Fig. 1**). The chromatographic separation is mainly disturbed by the high concentration of chlorophyll *a* and carotenoids, which are about 1/4 of the total acyl lipid materials by mass. We avoided this by applying the total lipid extract on the plate as a narrow band, leaving a blank space on the right for applying standards later. This way, as much as 5 – 15 mg of acyl lipids could be loaded on a single plate.

**Fig. 1.**
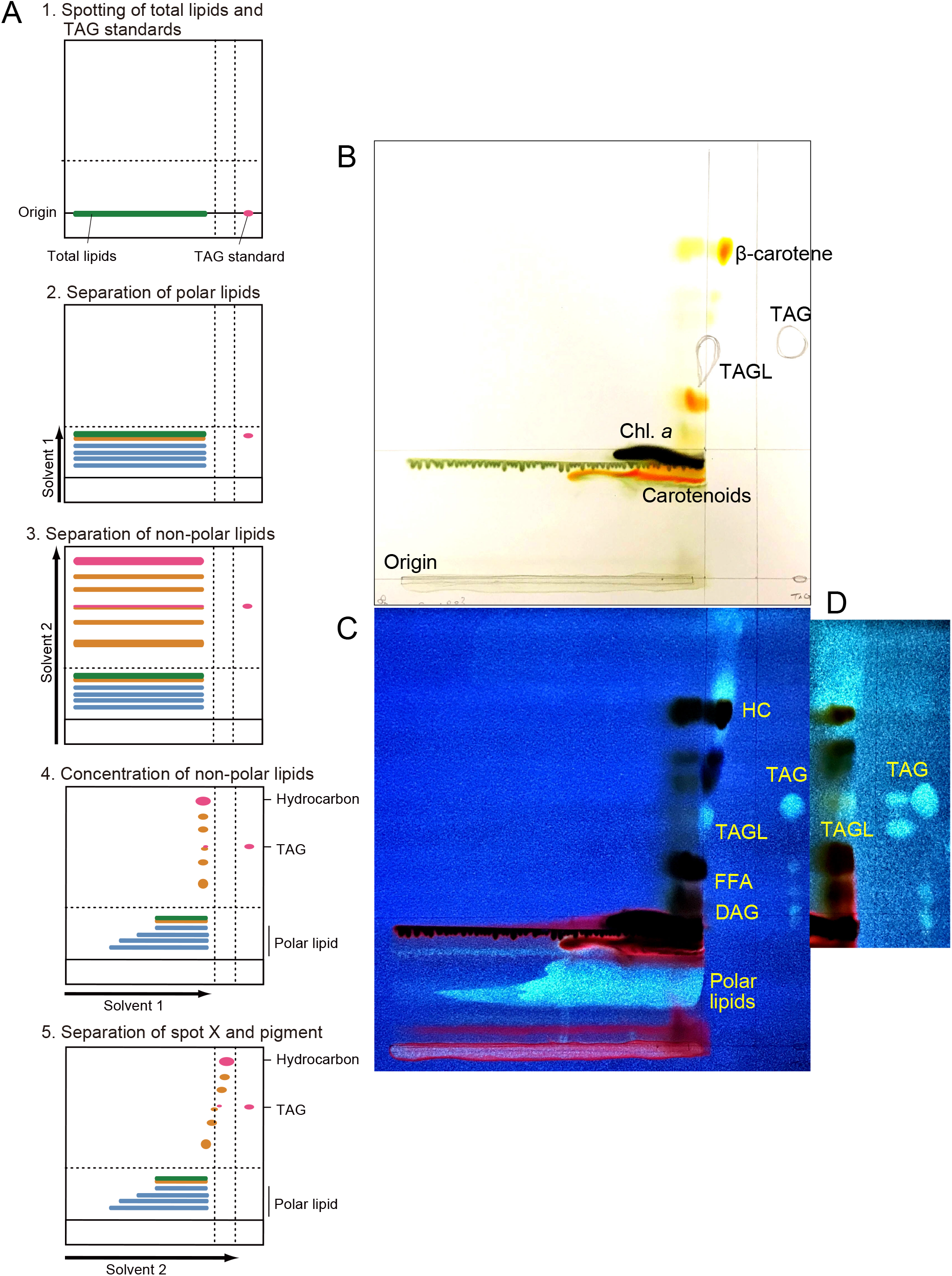
Isolation of TAGL fraction by two-dimensional TLC. (A) Explanation of development method. Solvents: 1, acetone/toluene/methanol/water (8:3:2:1, by volume); 2, *n*-hexane/diethyl ether/acetic acid (80:30:1, by volume). See the text for a description of the method. (B) Isolation of TAGL from *Synechocystis*. Spots of TAG and TAGL detected by fluorescence were marked by pencil. (C) Detection of lipids by the fluorescence of primuline for the plate in (B). (D) Fluorescence detection of TAG and TAGL in *Nostoc*. HC, hydrocarbons; FFA, free fatty acids; DAG, diacylglycerol. TAG standards: triolein in (B) and (C); triheptadecanoin (left) and triolein (right) in (D). Note that the TAG standards were partially degraded.

In the first dimension, a polar solvent mixture was used to develop all major glycerolipids to the middle of the plate. After complete drying *in vacuo*, the plate was further developed in the same direction with a non-polar solvent until the top for separating non-polar lipids such as hydrocarbons, TAG (if present), and free fatty acids, along with pigments. After another drying, the plate was developed with a polar solvent from left to right, leaving a narrow blank area. Finally, another development with a non-polar solvent displaced non-polar lipids to the narrow blank area before the standards area. This procedure achieved a concentration of non-polar lipids that had been applied in bands to single spots along the final front line.

We detected non-polar lipids on the final front line, such as diacylglycerol, free fatty acids, and hydrocarbons. We also detected a spot co-migrating authentic TAG standards. We named it TAG-like or TAGL and further analyzed it. The exact position of TAGL relative to TAG was somewhat variable depending on TLC development, most probably because the third and fourth developments were not precisely straight. However, we consistently detected a single spot of TAGL in *Synechocystis* sp. PCC 6803 (*Synechocystis*), *Anabaena* sp. PCC 7120, and *Nostoc punctiforme* PCC 73102 (*Nostoc*). We analyzed mainly the materials from *Synechocystis* and *Nostoc*.

### Analysis of TAGL

TAGL was purified from *Synechocystis* and *Nostoc* and subjected to methanolysis. The fatty acid methyl esters were analyzed (**Tables 1 and 2**). Major fatty acids were saturated fatty acids, such as palmitic (16:0) and stearic (18:0) acids, as reported previously (Paramuna and Sumners 2014). The levels of these fatty acids varied with samples, as indicated by the large standard deviation. The level of TAGL, as judged from the fatty acid content, was 0.41±0.11% and 0.32±0.06% in *Synechocystis* and *Nostoc*, respectively.

**Table 1.** Fatty acid composition of TAGL fraction in *Synechocystis*. Each value is a percentage of the total (average ± standard deviation).

| Fatty acid | Total lipids<br>($n = 4$) | TAGL<br>($n = 3$) |
| --- | --- | --- |
| 14:0 | 0.1±0.2 | 3.1±1.0 |
| 16:0 | 46.4±13.2 | 37.3±10.8 |
| 16:1(9) | 3.3±0.6 | 0.8±0.6 |
| 17 : 0 | 0.3±0.1 | 2.2±1.1 |
| 17:1(10) | 0.9±0.3 | 0.0±0.0 |
| 18:0 | 0.9±0.2 | 29.6±13.2 |
| 18:1(9) | 12.4±1.8 | 23.0±18.7 |
| 18:2(6,9) | 0.4±0.0 | 0.0±0.0 |
| 18:2(9,12) | 13.6±0.8 | 1.4±0.5 |
| 18:3(6,9,12) | 13.6±2.4 | 2.2±1.2 |
| 18:3(9,12,15) | 0.8±0.4 | 0.5±0.6 |
| 18:4(6,9,12,15) | 0.4±0.2 | 0.0±0.0 |

Fatty acids are shown by convention (carbon number : number of double bonds). Positions of double bonds are shown in parenthesis.

**Table 2.** Fatty acid composition of TAGL fraction in *Nostoc*. Each value is a percentage of the total (average ± standard deviation).

| Fatty acid | Total lipids<br>(*n* = 3) | TAGL<br>(*n* = 2) |
| --- | --- | --- |
| 14:0 | 0.1±0.1 | 4.6±6.5 |
| 16:0 | 29.6±1.5 | 27.8±6.2 |
| 16:1(9) | 16.9±1.5 | 4.4±5.8 |
| 16:2(9,12) | 0.7±0.0 | 0.0±1.1 |
| 18:0 | 2.9±0.3 | 50.5±20.0 |
| 18:1(9) | 8.6±0.8 | 4.8±2.1 |
| 18:1(11) | 1.1±0.1 | 0.7±0.9 |
| 18:2(9,12) | 35.9±0.6 | 5.4±1.5 |
| 18:3(9,12,15) | 3.7±0.9 | 0.5±0.1 |
| 20:0 | 0.2±0.1 | 0.0±0.0 |
| 20:2(11,14) | 0.3±0.2 | 0.0±0.0 |

Fatty acids are shown by convention (carbon number : number of double bonds). Positions of double bonds are shown in parenthesis.

TAGL was analyzed by MALDI TOF-MS to estimate the molecular weight (**Fig. 2**). Peaks were detected at about *m*/*Z*=1011 and 1039. Assuming these are Na^+^ adducts, their monoisotopic masses should be 988 and 1016, respectively. Assuming that these correspond to palmitoyl and stearoyl monoesters, the remaining alcohol part has a molecular weight of 750.

**Fig. 2.**
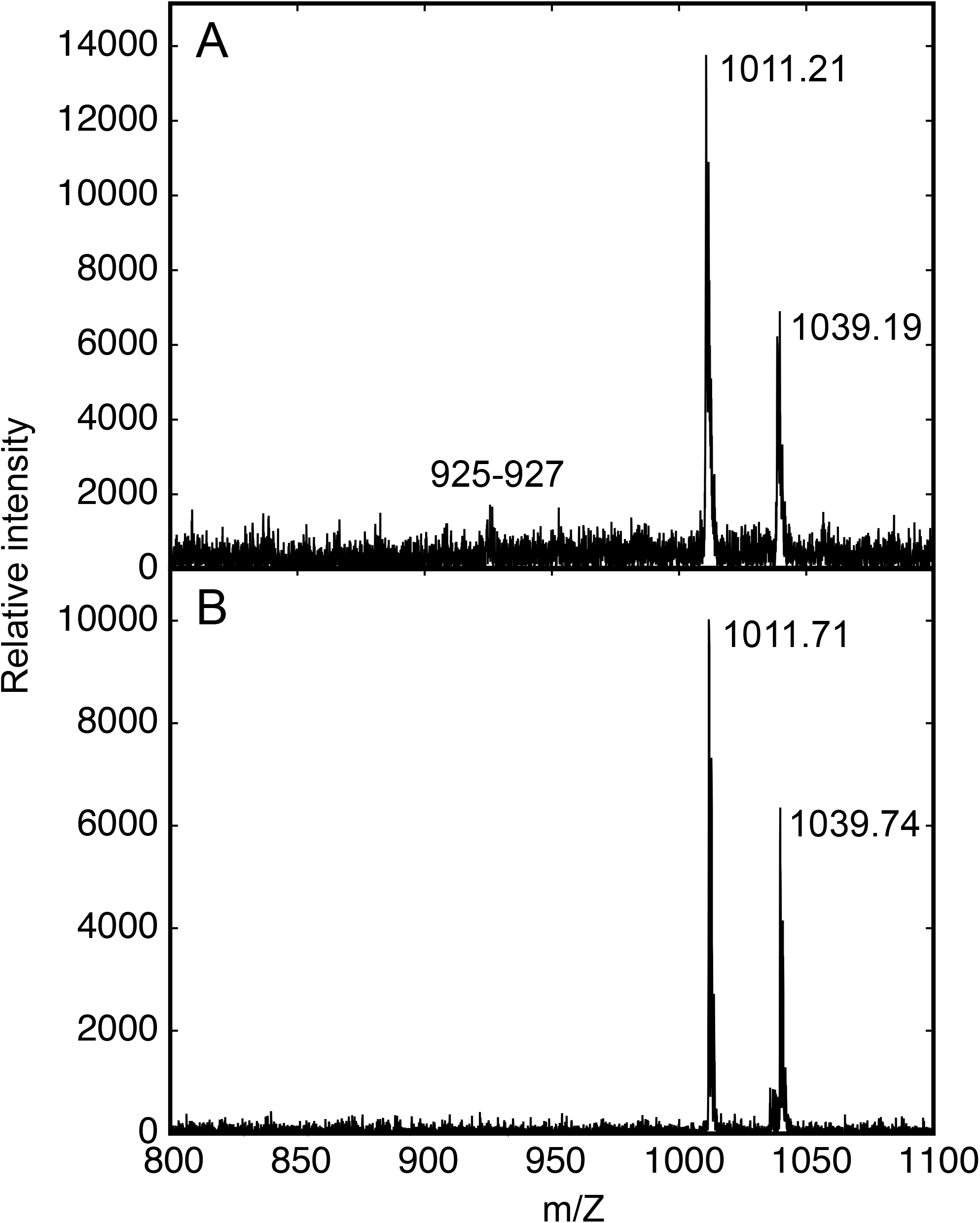
MALDI TOF-MS analysis of TAGL. (A) *Synechocystis* (B) *Nostoc*.

Peaks of TAG (as Na^+^ adduct) should appear in the region *m*/*Z*=800 – 1000: 48:0 (tripalmitin) at 829 and 54:0 (tristearin) at 913, which we confirmed by analysis with standards. TAG fraction from spinach chloroplast indeed showed peaks at 909 (54:2), 907 (54:3), 885 (52:0), and 857 (50:0) (results not shown). Since TAG usually gives high-intensity signals in MALDI TOF-MS, the level of TAG, if present, must be minimal (noise level) in cyanobacteria.

### NMR analysis

^1^H- and ^13^C-NMR analysis gave interesting clues to the elucidation of the structure of TAGL. The ^13^C-NMR spectrum (**Fig. 3**) showed that TAGL is a monoester (signal at 172.7 ppm). Other signals were found in two regions, 10 – 40 ppm and 120 – 150 ppm. No signals that could be assigned to the glycerol moiety of TAG (typically, at 69 and 62 ppm. Gunstone 1994) were detected. The spectrum suggested a simple structure consisting of many repeating units.

**Fig. 3.**
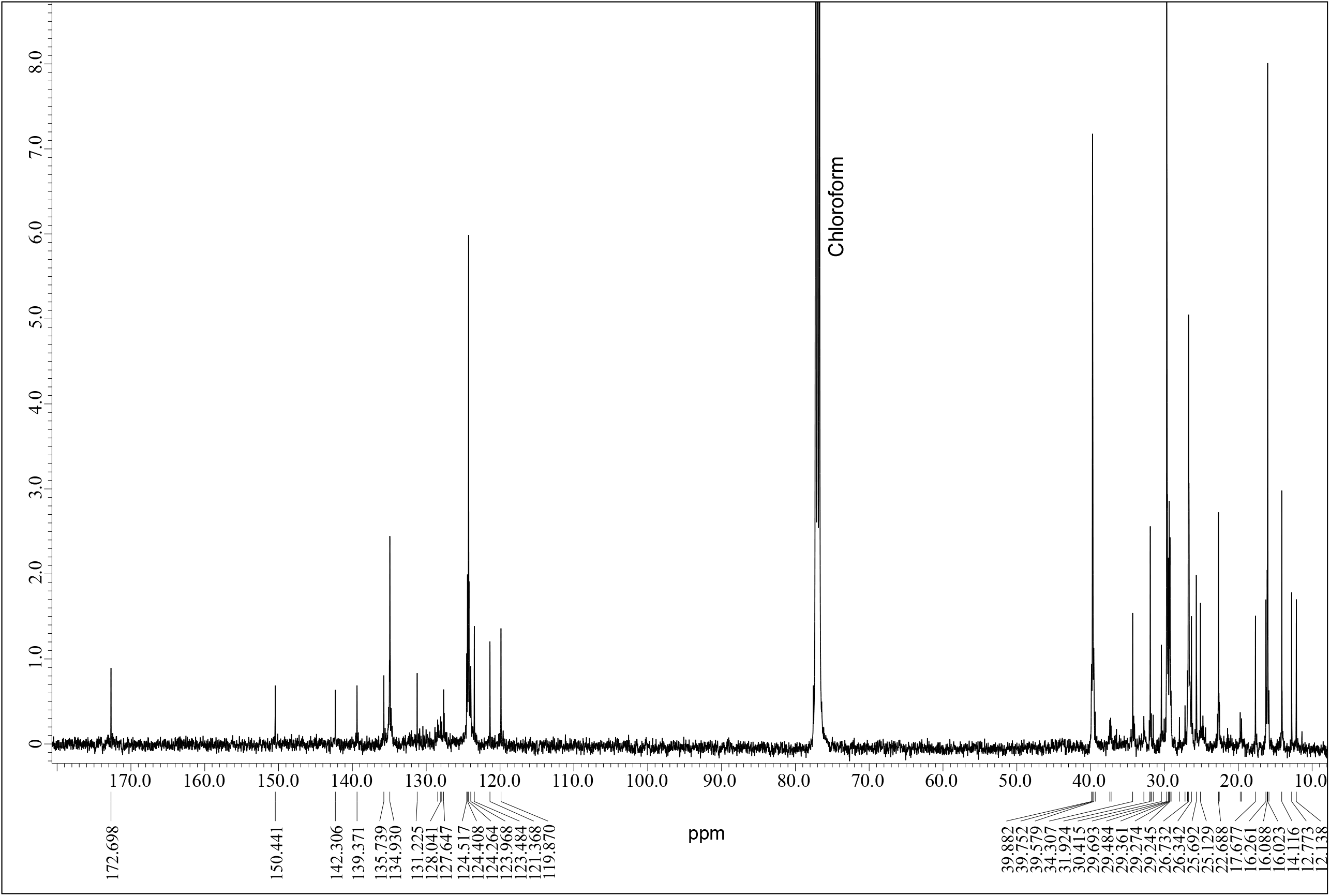
^13^C NMR spectrum of TAGL isolated from the *Synechocystis* cells grown with partial ^13^C supplementation. Tetramethyl silane was used as the reference for the chemical shift. Note that the large peak at 77 ppm represents chloroform (solvent). For the detail of the peak assignment, see Table 3.

The results of ^1^H-NMR were more complicated (**Fig. 4**). The characteristic peaks at 3.3 ppm (doublet) and 6.6 ppm (singlet) were not found in available NMR databases. Two dimensional NMR analysis called COSY (**Supplementary Fig. S1**) suggested a coupling of 3.3 ppm and 5.3 ppm for double bonds (green circles). A weak coupling of the 3.3 ppm peak and the 6.6 ppm peak was also found, interpreted as a long-range coupling. The 3.3 ppm doublet should arise from CH_2_ adjacent to a double bond, and the 6.6 ppm signal could be an aromatic hydrogen, and they are linked. This particular structure is rarely found in the literature but could be ascribed to plastoquinone. No NMR spectrum of plastoquinone is available in the current databases. However, we found that the initial studies of plastoquinone discovery already described 60 MHz ^1^H NMR data, including the 3.3 and 6.6 ppm signals being the decisive evidence for the structure (Eck and Trebst 1963).

**Fig. 4.**
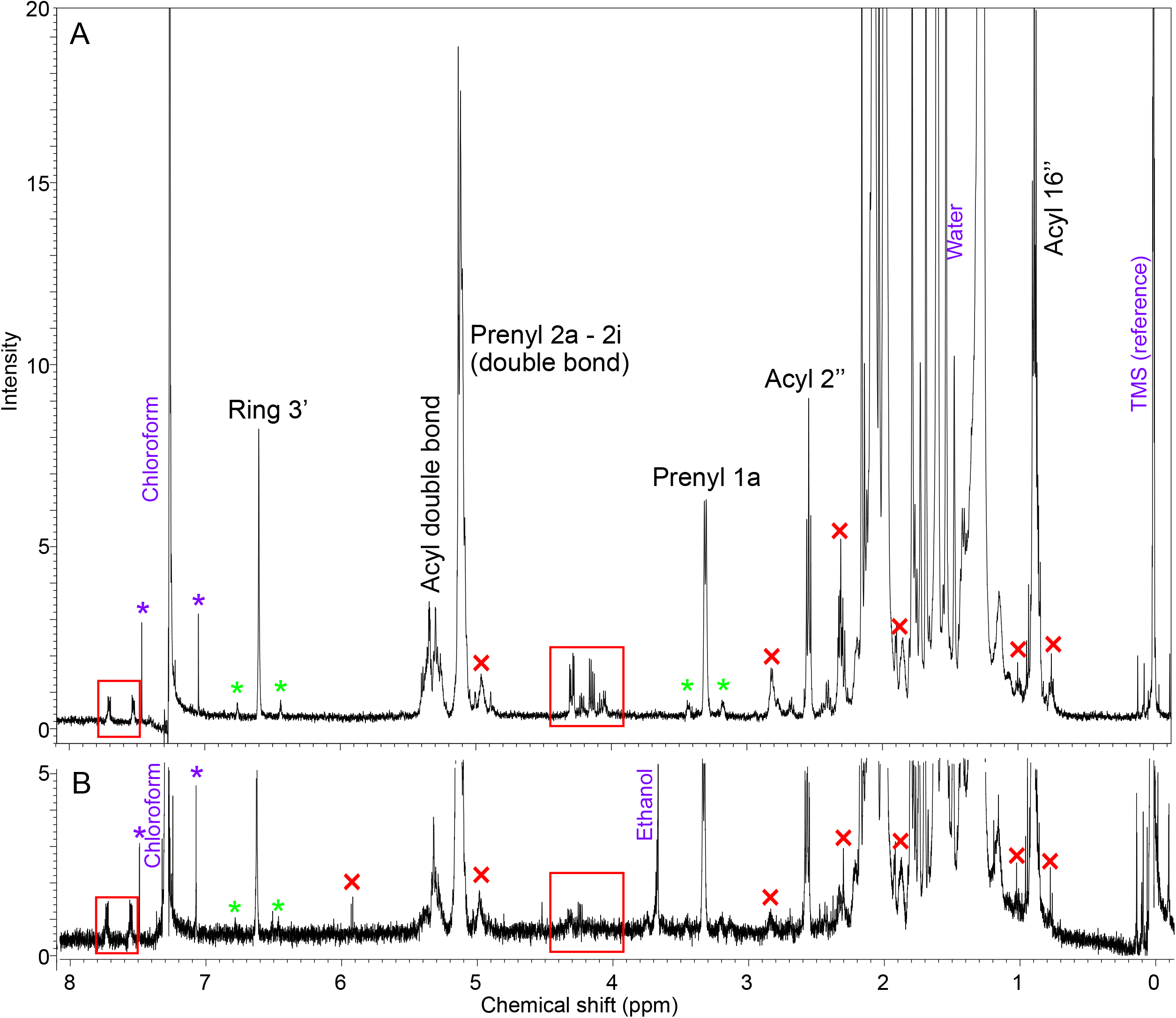
^1^H NMR spectra of two preparations (A and B) of TAGL from the *Synechocystis* cells grown with partial ^13^C supplementation. Tetramethyl silane (TMS) was used as the reference for the chemical shift. Only small peaks are shown in spectrum B. Note that the two regions marked by rectangles and the peaks marked by crosses represent contaminants or minor components (see Table 4). The asterisks indicate satellite peaks arising from ^13^C-^1^H coupling (green asterisks for TAGL and purple asterisks for chloroform). The solvent chloroform appears at 7.2 ppm. The large peak at 1.5 ppm is due to water. For the detail of the peak assignment, see Table 3. For atom numbers, see Fig. 5.

As described above, the mass spectral analysis suggested that the alcoholic residue had a molecular weight of 750. This value corresponds to the monomolecular weight of plastoquinol-9 (a reduced form of plastoquinone-9). Therefore, we considered acyl plastoquinol (APQ) as a candidate for TAGL (**Fig. 5**). Because no reference compound was available, we used some web services of NMR simulation (http://www.nmrdb.org and https://nmrshiftdb.nmr.uni-koeln.de) to test the possibility that the structure of acyl plastoquinol is compatible with the observed NMR signals (**Supplementary table S1**). Note that two isomers are possible for APQ, depending on the position of the acyl group. The simulation did not give differences in most parts of the molecule in the two possible isomers (both ^1^H and ^13^C), and the results were consistent with the observed data. We suspect that the 1-*O*-acyl isomer fits better with the observed data, but this is not decisive. Accordingly, we could assign all carbons and hydrogens as summarized in **Table 3**. Currently, we still need to determine the position of acylation.

**Fig. 5.**
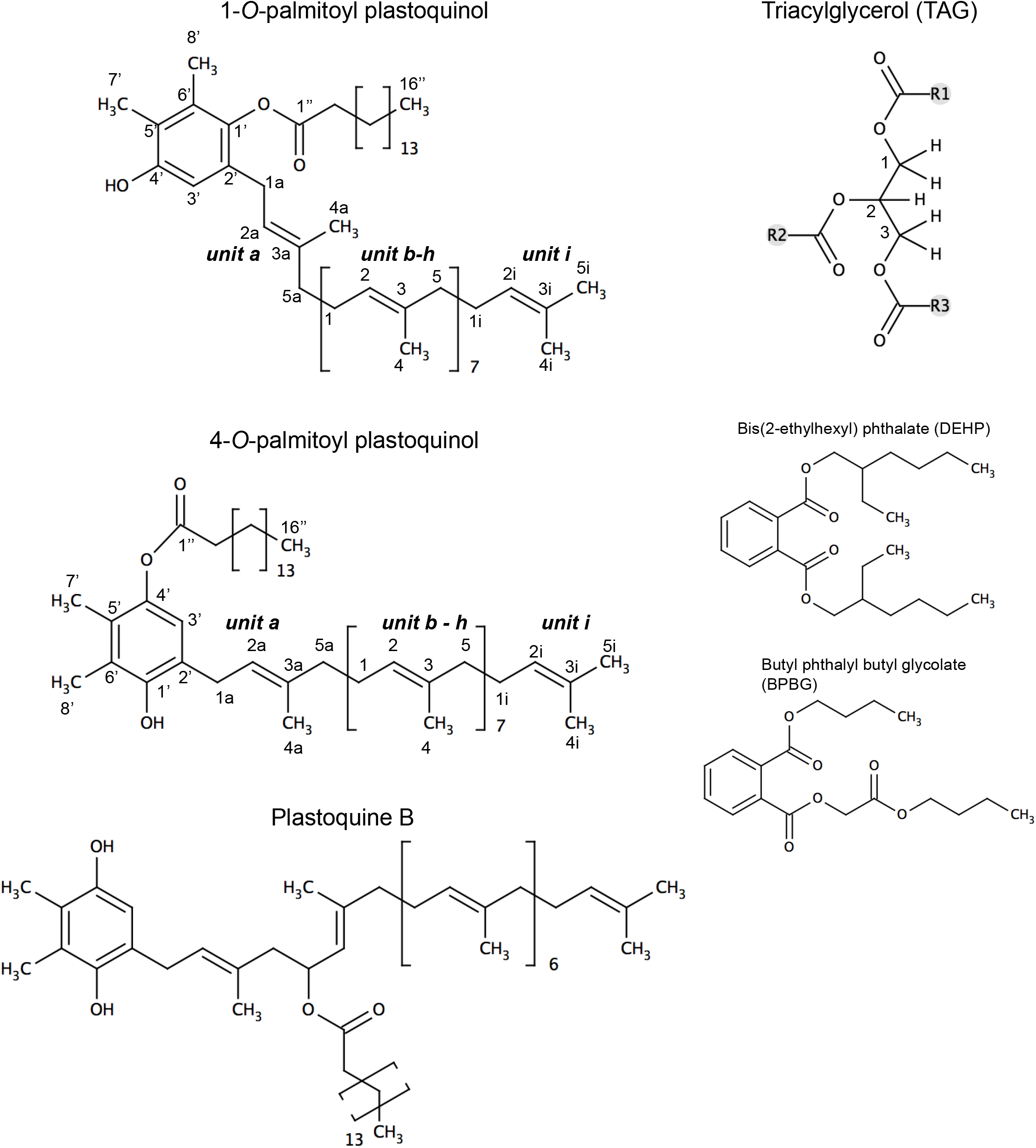
Structures of APQ (two isomers), plastoquinone B, triacylglycerol, and two possible contaminants (phthalate esters). In APQ, carbon atoms are numbered. In the prenyl chain, the five carbons in each C5 unit are numbered from 1 to 5, with an alphabet representing each unit. Carbons in the aromatic ring are numbered with a prime. Carbons in the acyl chain are numbered with a double prime.

**Table 3.**
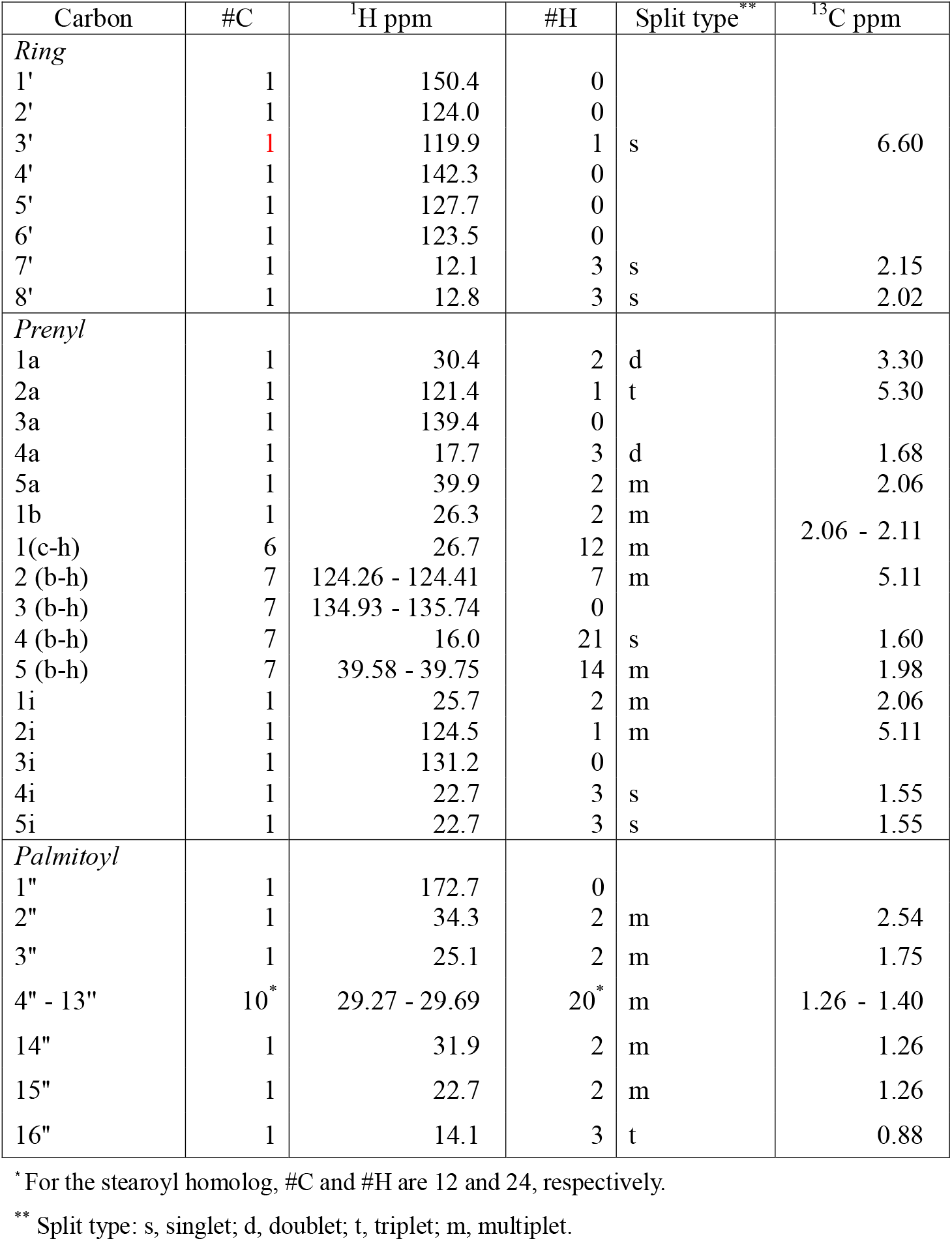
Assignment of NMR signals of palmitoyl plastoquinol. #C and #H represent the number of carbons and hydrogens, respectively.

### Minor peaks in NMR

However, we found various small signals in the ^1^H-NMR spectra (**Table 4** and **Fig. 4**). We analyzed two preparations of TAGL by NMR, but the minor signals were not identical. The intensity of some signals was also different in the two preparations (compare panels A and B in **Fig. 4**). The variable signals might not be authentic signals of the major component of TAGL. The region 4.1 – 4.5 ppm (red rectangle) could represent the glycerol moiety of TAG. The quartets at 4.29 and 4.15 ppm were coupled to each other and also coupled to ^13^C at 62.2 and 65.7 ppm, which were not visible in the spectrum in **Fig. 3**, but detected by HSQC analysis (**Supplementary Fig. S2**). The 65.7 ppm peak was somewhat different from the typical value for TAG glycerol (69 ppm). These carbons were not enriched with ^13^C during the growth and could be contaminants. Alternatively, TAG synthesis could be very slow to be labeled effectively. Another problem in assigning these signals to TAG was the absence of C2 hydrogen (typically at 5.3 ppm) (**Fig. 4**) which should appear in the COSY spectrum (**Supplementary Fig. S1**).

**Table 4.** Minor component peaks in ^1^H-NMR. #H shows the peak area relative to the single hydrogen of APQ at 6.6 ppm.

| Peak (ppm) | #H | COSY | HSQC | Tentative assignment |
| --- | --- | --- | --- | --- |
| 7.70 | 0.10 – 0.17 | 7.53 | 128.4 | Phthalate esters |
| 7.53 | 0.10 – 0.14 | 7.70 | 131.2 | Phthalate esters |
| 4.96 | 0.5 | -- | 111.0 | Unknown double bond |
| 4.89 | 0.7 | -- | 111.0 | Unknown double bond |
| 4.29 | 0.14 – 0.30 | 4.15, 5.30 | 62.2, 65.7 | Phthalate esters (BPBG?),<br>glycerol of TAG |
| 4.22 | 0.10 – 0.18 | 1.72 | -- | Phthalate esters (DEHP?) |
| 4.15 | 0 – 0.20 | 4.29, 5.30 | 62.2 | Phthalate esters (BPBG?),<br>glycerol of TAG |
| 4.09 | 0 – 0.10 | 1.55 | -- | Unknown |
| 2.82 | 0.22 – 0.50 | 1.26, 5.35 | -- | Acyl -CH=CH-CH<sub>2</sub>-CH=CH- |
| 2.31 | 1.0 | 1.60 | 34.3 | Acyl 2" (alkyl ester) |
| 1.86 | 0.60 | -- | 43.7 | Unknown |
| 0.92 | ca. 1.0 | -- | -- | Phthalate esters, linoneyl<br>methyl |
| 0.86 | ca. 1.0 | -- | -- | Unknown (methyl) |
| 0.75 | 0.10 | 1.60 | 14.1 | Unknown (methyl) |

The small but significant peaks at 7.70 and 7.53 ppm are typically ascribed to aromatic hydrogens. Analysis of coupling (COSY and HSQC) also suggested the aromatic origin of these signals. Phthalate esters, common plasticizers, could explain these signals. We were not able to pinpoint the compound(s), but bis(2-ethylhexyl) phthalate (DEHP), which is the most used phthalate plasticizer, was a candidate. DEHP could also be the origin of the significant signal at 4.22 ppm in panel B (**Fig. 4**). Another commonly used phthalate, butyl phthalyl butyl glycolate (BPBG), could be an origin of signals at 7.7, 7.5, 4.3, and 4.1 ppm. Phthalate esters also present methyl signals at about 0.95 ppm, which could explain the 0.92 ppm signal. Although other phthalate signals were not identified among the large and crowded signals in **Fig. 4**, phthalate esters are likely contaminants of TAGL.

Other peaks at 2.82 ppm and 2.31 ppm were assigned to the unsaturated acyl groups (Vigli et al. 2003) that were present in TAGL preparation as minor components (**Table 1**). We could not explain all minor peaks, but they cannot be ascribed to any TAG part.

### Estimation of the TAG level

The TAG level was below the noise level in both MALDI TOF-MS and NMR analyses. However, we cannot exclude the presence of TAG in the TAGL preparations. We may estimate the maximum level of TAG that could be present in the cyanobacteria analyzed. If the 4.29 and 4.15 ppm signals in ^1^H-NMR were entirely due to TAG glycerol (two hydrogens each for C1 and C3), the total intensity must be less than 40% of the 6.6 ppm signal (single hydrogen in APQ). Dividing by four, the maximum TAG level must be 10% of APQ (mole basis). As described above, not all TAG signals could be detected, and phthalate esters could contribute to the C1 and C3 signals, the level of TAG must be much lower.

### Lipid droplets in cyanobacteria

Microscopic examination showed cyanobacterial cells contained tiny lipid droplets that fluoresce with BODIPY staining (**Fig. 6A-D**). Transmission electron micrographs showed small electron-dense particles corresponding to the fluorescent lipid droplets (**Fig. 6E-H**). Previous works suggested that cyanobacterial lipid droplets contain hydrocarbons and, possibly, TAG (Peramuna and Summers 2014, Peramuna et al. 2015). The level of hydrocarbon (mainly heptadecane) was 59.7±27.4 and 98.3±22.3 nmol µmol^-1^ fatty acids in *Synechocystis* and *Nostoc*, respectively, or about 6 – 10% of the total fatty acids. According to the results, alkanes must be the major component of lipid droplets in cyanobacteria. The contribution of APQ must be quite limited.

**Fig. 6.**
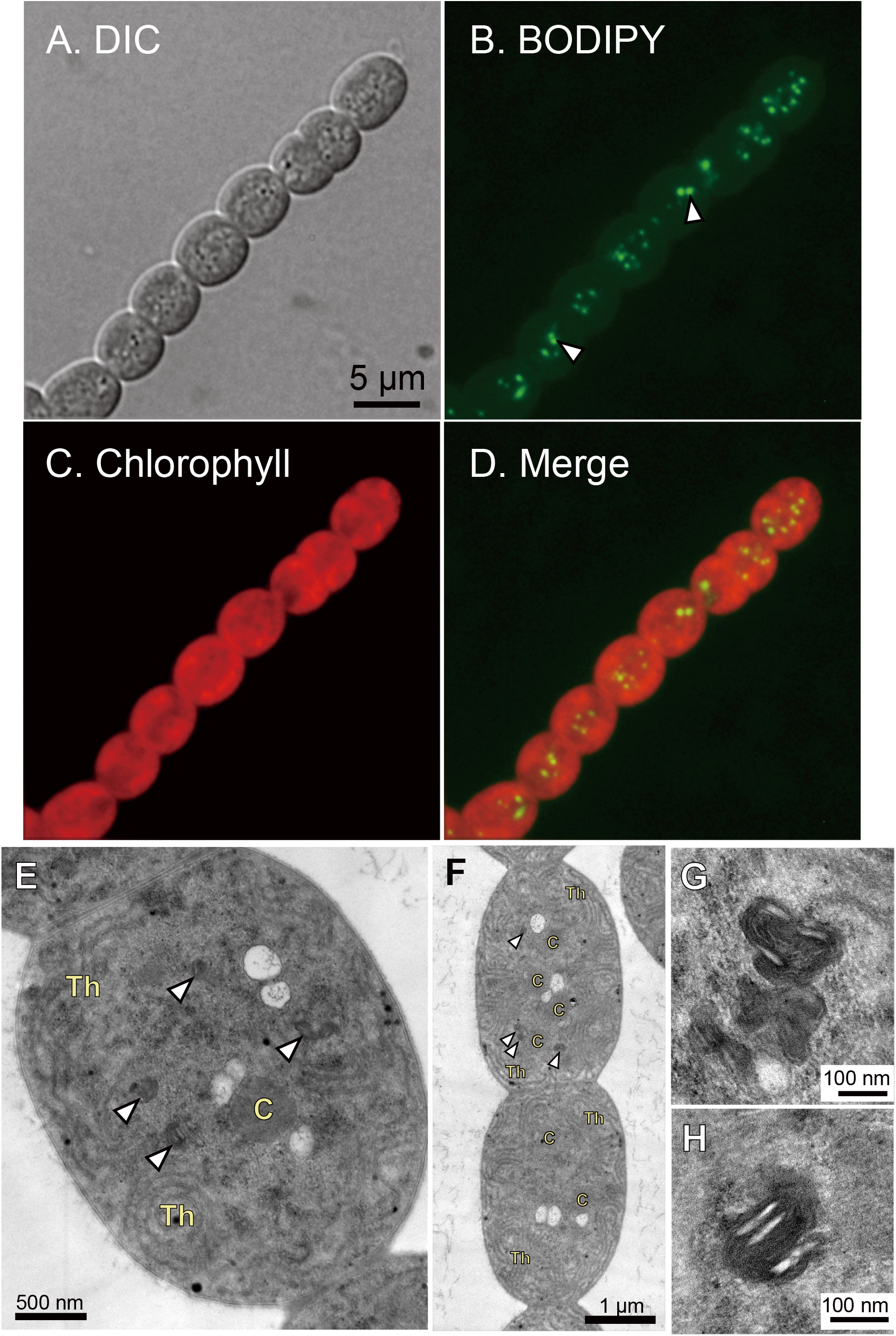
Lipid droplets in *Nostoc* cells. (A) – (D), fluorescence micrographs after BODIPY staining. DIC, differential interference. (E) – (H), Transmission electron micrographs. (G) and (H) show enlarged views of lipid droplets. Each arrowhead indicates a lipid droplet. C, carboxysome; Th, thylakoid membrane.

## Discussion

We found acyl plastoquinol (APQ) in cyanobacteria. Plastoquinone is an essential component in photosynthesis, and its synthesis in cyanobacteria has been established (Nowicka and Kruk 2016). APQ co-migrates with TAG in a standard TLC system used to analyze TAG, which is why many previous studies believed APQ was TAG. Nevertheless, we cannot exclude the possibility that TAG exists in cyanobacteria at a low level. We estimate that the level of APQ is about 0.4% of the total lipid fatty acids, and the TAG level, if present, must be ten times lower or much less (<< 0.04% of the total lipid fatty acids). In this respect, the reported TAG levels were too high. Aizouq et al. (2020) described “5 nmol TAG OD750^-1^” and “4.9 µg Chl OD760^-1^” (5.5 nmol Chl OD760^-1^). These values are similar to each other if the volume basis is identical (theoretically, 1 mL), which must be an error. Assuming that the TAG value was expressed for a 100 mL culture, the TAG level should be 0.05 nmol OD750^-1^ mL^-1^, roughly 1% of chlorophyll. Assuming that the chlorophyll content is about 1/8 of the total lipid fatty acids (typical value of our analysis), the TAG level was 0.4% of the total lipid fatty acids. In this calculation, TAG level is expressed in mol TAG basis, not TAG fatty acids. The level reported by Tanaka et al. (2020) was 0.01 – 0.03 nmol TAG mL^-1^ OD730^-1^, equivalent to 0.08 – 0.24% of the total lipid fatty acids (or somewhat more because OD730 > OD750 > 0D760). This level was also higher than our estimated maximum value. Conversely, we could suspect that the TAGL spot did not precisely overlap TAG in the TLC system, and our TAGL samples included only part of TAG. However, the reported fatty acid composition or molecular species composition of TAG (both Aizouq et al. 2020 and Tanaka et al. 2020) included a significant level of polyunsaturated fatty acids, which was not consistent with other previous reports using one-dimensional TLC in which TAG and TAGL co-migrates. Although Tanaka et al. (2020) discussed that the detected TAG was not due to contamination, we still have to ask why liquid chromatography used by Aizouq et al. (2020) and Tanaka et al. (2020) gave higher TAG values. The minute level of TAG that we estimated could be synthesized by various acyl transfer reactions, either enzymatic or non-enzymatic. The inability of ^13^C labeling of TAG is compatible with the non-enzymatic synthesis or contamination. The ultimate identification of cyanobacterial TAG must be a complex problem requiring sophisticated means.

Plastoquinone B, or PQB, has been known as an acylated plastoquinone (Das et al. 1967). It is a hydroxylated plastoquinone (oxidized form), esterified with an acyl group. PQB was described just after the discovery of plastoquinone (plastoquinone A at the time). PQB bearing a palmitoyl group was found to have *m*/*Z*=1002, which corresponds to acyloxy plastoquinone, having a hydroxyl group at the methylene carbon between two double bonds (**Fig. 5**). The structure of PQB is sometimes erroneously described as acylated hydrated plastoquinone, which must have *m*/*Z*=1004. In any case, APQ is different from PQB. PQB is a hydroxylated plastoquinone acylated at the added hydroxyl group. It does not become plastoquinone by de-acylation. APQ, on the other hand, releases plastoquinol if hydrolyzed. Alternatively, it becomes plastoquinone (oxidized form) upon oxidation by releasing a fatty acid. In this respect, although the amount is small, APQ could be a reservoir of reducing power within the cell. Another function of APQ could be the protection of photosystem II, or it helps the regeneration of photosystem II after photoinhibition.

A remaining question is whether APQ is present in only cyanobacteria. If the Slr2103 protein synthesizes APQ, this enzyme is not present in all cyanobacteria. *Synechococcus* and *Prochlorococcus* do not have a homolog of Slr2103. Phylogenetic analysis indicates that cyanobacterial Slr2103 homologs and actinobacterial counterparts are a sister group of plant PES1 (**Supplementary Fig. S3**). The tree is a revised version of the phylogenetic tree previously shown by Aizouq et al. (2020). A detailed analysis of the LPAAT part was previously published (Sato and Awai 2017). We now confirm the sister relationship of Slr2103 and PES1, which suggests that plant PES1 could also synthesize APQ within the chloroplast. However, the structure of PES1 is very complex, with additional domains, which could limit substrate specificity.

## Materials and Methods

### Growth of cells

*Synechocystis* sp. PCC 6803 strain GT-S (Tajima et al. 2012) and *Nostoc punctiforme* PCC 73102 were grown as described previously (Sato 2015). To prepare materials for ^13^C-NMR, [^13^C]-labeled sodium bicarbonate (Cambridge Isotope Laboratories, Inc.: 99% ^13^C) was added to the culture.

### Fluorescence microscopy

Lipid droplets were stained with BODIPY and examined under a fluorescence microscope, as described previously (Moriyama et al. 2018).

### Transmission electron microscopy

*Nostoc* cells were processed according to the “hot osmium” method, and the sections were examined by JEOL 1200EX operated at 100 kV (Sato et al. 2017).

### Lipid analysis

Total lipids were extracted from the harvested cells by Bligh and Dyer (1959) method. The total lipids from the 1.5 L culture were concentrated, dissolved in 2.5 mL of chloroform/methanol/ethanol (2:1:3, v/v/v), and stored at -16°C until analysis.

The total lipids were fractionated by two-dimensional TLC. Because the TLC analysis is an essential point in the current study, important points of the analysis are also described in **Results**. Here, additional methodological details are explained. Silica gel 60 plates (Merck Catalog No. 1.05721.0001) were initially developed with HPLC-grade methanol (FUJIFILM Wako Pure Chemicals, Osaka) to the top of the plate to eliminate any lipid materials that could have been adsorbed on the plate. It is essential to remove fatty acid-containing materials that could interfere with quantifying a minute amount of acyl-lipid before the main separation step. We used a vacuum desiccator to completely dry the plate after each development to ensure another development without being affected by the previous development with a different solvent mixture. Then, the total lipid extract was applied on the plate as a narrow band, leaving a blank space for standards on the right (**Fig. 1**). Typically, as much as 5 − 15 mg acyl lipids (equivalent to 1.5 – 4.5 mg chl. *a*) could be loaded on a single plate. The chromatographic development is mainly disturbed by the high concentration of chlorophyll *a* and carotenoids, which are about 1/4 of the total acyl lipid materials. In the current scale of analysis, the total lipids were cleanly and safely separated without serious disturbance by the pigments.

In the first dimension (upwards in **Fig. 1**), acetone/toluene/methanol/water (8:3:2:1, by vol.) was used to develop all major glycerolipids to the middle of the plate. After complete drying *in vacuo*, the plate was further developed in the same direction with *n*-hexane/diethyl ether/acetic acid (80:30:1, by vol.) to the top of the plate to separate non-polar lipids such as hydrocarbons, TAG (if present), and free fatty acids, along with pigments. After drying *in vacuo* again, the plate was developed with acetone/toluene/methanol/water (8:3:2:1, by vol.) from left to right, leaving a blank area before the standards. Finally, development with *n*-hexane/diethyl ether/acetic acid (80:30:1, by vol.) displaced non-polar lipids further. This development compressed the bands of non-polar lipids to single spots aligning on the front line. The spot (TAGL) co-migrating with authentic triolein or triheptadecanoin was recovered from the silica gel and stored in chloroform/methanol (2:1, v/v) at -16°C.

Fatty acids were analyzed according to the established method in the laboratory (Sakurai et al. 2014). Alkanes were recovered from the “HC” spot of the TLC plates (**Fig. 1**) and analyzed by GC/MS using eicosane as an internal standard.

### MALDI TOF-MS analysis

TAGL fraction was recovered from the plate and analyzed by MALDI TOF-MS using 2,5-dihydroxybenzoic acid as the matrix. Mass spectra were recorded in an AXIMA-CFR Plus matrix-assisted laser desorption ionization (MALDI)–time of flight (TOF) mass spectrometer (Shimadzu, Kyoto, Japan) at an irradiation intensity corresponding to the setting at about 115. The GNU plot software was used to process the data for presentation.

### NMR analysis

^1^H-NMR (500 MHz) and ^13^C-NMR (125 MHz) were measured using a Bruker AVANCE III 500 spectrometer, using the TAGL materials purified from the culture with ^13^C supplementation and dissolved in CDCl_3_. GC/MS analysis of fatty acid methyl esters showed that about 10% of carbon was replaced with ^13^C. This level of ^13^C did not affect the measurement of ^1^H-NMR, although some satellite peaks due to ^13^C-^1^H coupling were visible.

The same machine was used for COSY and HSQC analyses.

### Phylogenetic analysis

Slr2103 homologs were collected as clusters using the Gclust2021 dataset in the Gclust database (Sato 2009: http://gclust.c.u-tokyo.ac.jp/). The phylogenetic trees were constructed by the Bayesian Inference method according to the procedure described in Sato and Awai (2017) and Sato (2020).

### Data Availability Statement

All data are presented in the paper. No new sequence data are presented.

## Supporting information

Supplementary Table 1 and three figures.

## Abbreviations

APQ: acyl plastoquinol
COSY: correlation spectroscopy
GC/MS: gas chromatography/mass spectrometry
HSQC: heteronuclear single quantum correlation
MALDI TOF-MS: matrix-assisted laser desorption ionization time-of-flight mass spectrometry
NMR: nuclear magnetic resonance spectroscopy
TAG: triacylglycerol
TAGL: triacylglycerol-like lipid
TLC: thin-layer chromatography.

## Acknowledgements

Phylogenetic calculations were performed in the supercomputer system of the Human Genome Center, the University of Tokyo. We are grateful for helpful discussions with Dr. Haruhiko Jimbo, University of Tokyo.

## Author Contributions

N.S. designed the research, N.M., T. Y., and N.S. performed the research, and N.S. wrote the paper.

