## Supplementary Table 1 and three figures. for "Acyl plastoquinol is a major substance that co-migrates with triacylglycerol in cyanobacteria"

Index

**Supplementary Table S1** Summary of NMR simulations
**Supplementary Fig. S1** COSY analysis of *Synechocystis* TAGL
**Supplementary Fig. S2** HSQC analysis of *Synechocystis* TAGL
**Supplementary Fig. S3** Bayesian Inference (BI) phylogenetic tree of acyltransferases

| Supplementary table S1. Summary of NMR simulations. | | | | | | | | | | | | |
|---|---|---|---|---|---|---|---|---|---|---|---|---|
| Carbon | | C shift (ppm) | | | | H shift (ppm) | | | | | | |
| | | nmrshiftdb 1-O-acyl | nmrshiftdb 4-O-acyl | #C | Obs | nmrdb 1-O-acyl | nmrdb 4-O-acyl | nmrshiftdb 1-O-acyl | nmrshiftdb 4-O-acyl | split type | #H | Obs |
| Ring | | | | | | | | | | | | |
| 1' | (OH) | 154.95 | 156.40 | 1 | 150.4 | none | none | none | none | | | |
| 2' | (prenyl) | 122.97 | 122.97 | 1 | 124.0 | none | none | none | none | | | |
| 3' | | 110.10 | 106.17 | 1 | 119.9 | 6.42 | 6.13 | 6.71 | 6.71 | 1 | 1 | 6.60 |
| 4' | (OH) | 146.90 | 142.30 | 1 | 142.3 | none | none | none | none | | | |
| 5' | (methyl) | 125.10 | 126.90 | 1 | 127.7 | none | none | none | none | | | |
| 6' | (methyl) | 120.08 | 121.30 | 1 | 123.5 | none | none | none | none | | | |
| 7' | methyl | 11.90 | 12.20 | 1 | 12.1 | 2.18 | 2.15 | 2.15 | 2.15 | 3 s | 3 | 2.15 |
| 8' | methyl | 11.06 | 12.60 | 1 | 12.8 | 2.19 | 2.21 | 2.15 | 2.15 | 3 | 3 | 2.02 |
| Prenyl | | | | | | | | | | | | |
| 1a | | 28.90 | 28.90 | 1 | 30.4 | 3.26 | 3.18 | 3.37 | 3.37 | 2 d | 2 | 3.30 |
| 2a | double | 121.55 | 121.55 | 1 | 121.4 | 5.33 | 5.32 | 5.22 | 5.22 | 1 t | 1 | 5.30 |
| 3a | double | 140.00 | 140.00 | 1 | 139.4 | none | none | none | none | | | |
| 4a | methyl | 16.70 | 16.70 | 1 | 17.7 | 1.60 | 1.60 | 1.56 | 1.56 | 3 d | 3 | 1.68 |
| 5a | | 39.66 | 39.66 | 1 | 39.9 | 2.12 | 2.11 | 2.08 | 2.08 | 2 m | 2 | 2.06 |
| 1b | | 26.41 | 26.41 | 1 | 26.3 | 2.23 | 2.23 | 2.11 | 2.11 | 2 m | 2 | 2.06, 2.11 |
| 1(c-h) | | 26.60 | 26.60 | 6 | 26.7 | 2.21 | 2.21 | 2.11 | 2.11 | 2 m | 12 | |
| 2 (b-h) | double | 124.27 | 124.27 | 7 | 124.26 - 124.41 | 5.29 | 5.29 | 5.11 | 5.11 | 1 m | 7 | 5.11 |
| 3 (b-h) | double | 135.04 | 135.04 | 7 | 134.93 - 135.74 | none | none | none | none | | | |
| 4 (b-h) | methyl | 15.90 | 15.90 | 7 | 16.0 | 1.57 | 1.57 | 1.60 | 1.60 | 3 s | 21 | 1.60 |
| 5 (b-h) | | 39.70 | 39.70 | 7 | 39.58 - 39.75 | 2.10 | 2.11 | 1.98 | 1.98 | 2 m | 14 | 1.98 |
| 1i | | 26.70 | 26.70 | 1 | 25.7 | 2.20 | 2.20 | 2.08 | 2.08 | 2 m | 2 | 2.06 |
| 2i | double | 124.00 | 124.00 | 1 | 124.5 | 5.24 | 5.24 | 5.09 | 5.09 | 1 m | 1 | 5.11 |
| 3i | double | 131.79 | 131.79 | 1 | 131.2 | none | none | none | none | | | |
| 4i | methyl (cis) | 17.73 | 17.73 | 1 | 22.7 | 1.54 | 1.54 | 1.61 | 1.61 | 3 s | 3 | 1.55 |
| 5i | methyl (trans) | 25.76 | 25.76 | 1 | 22.7 | 1.54 | 1.54 | 1.67 | 1.67 | 3 s | 3 | 1.55 |
| Palmitoyl or stearoyl | | | | | | | | | | | | |
| 1'' | carboxyl | 172.26 | 172.26 | 1 | 172.7 | none | none | none | none | | | |
| 2'' | | 34.23 | 34.23 | 1 | 34.3 | 2.41 | 2.41 | 2.59 | 2.59 | 2 | 2 | 2.54 |
| 3'' | | 25.19 | 25.19 | 1 | 25.1 | 1.56 | 1.56 | 1.57 | 1.57 | 2 | 2 | 1.75 |
| 4'' - 13'' | | 29.35 - 29.68 | 29.35 - 29.68 | 10 - 11 | 29.27 - 29.69 | 1.229 - 1.264 | 1.229 - 1.263 | 1.20 - 1.25 | 1.20 - 1.25 | 2 | 20 - 24 | 1.26, 1.40 |
| 14'' | | 32.02 | 32.02 | 1 | 31.9 | 1.24 | 1.24 | 1.24 | 1.24 | 2 | 2 | 1.26 |
| 15'' | | 22.71 | 22.71 | 1 | 22.7 | 1.28 | 1.28 | 1.28 | 1.28 | 2 | 2 | 1.26 |
| 16'' | methyl | 14.10 | 14.10 | 1 | 14.1 | 0.87 | 0.87 | 0.87 | 0.87 | 3 | 3 | 0.88 |
| | | | | | | http://www.nmrdb.org | | https://nmrshiftdb.nmr.uni-koeln.de | | | | |

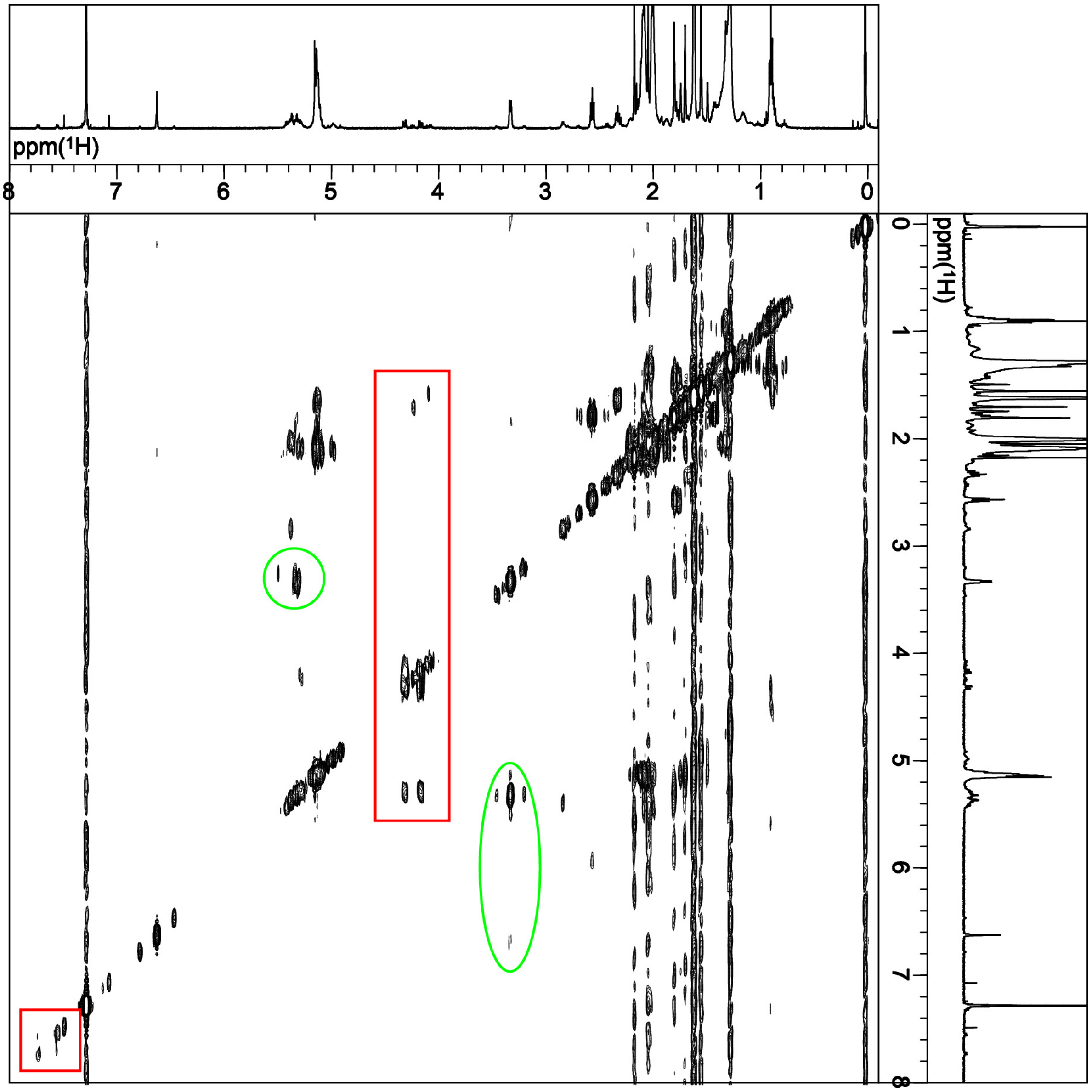


**Supplementary Fig. S1** COSY analysis of *Synechocystis* TAGL. A coupling of proximal $^1$H nuclei is detected. The signals characteristic of plastoquinonol structure are marked with green circles and ellipses. They are prenyl 1a (3.30 ppm) and ring 3' (6.60 ppm). A long-range coupling between these two atoms is found (lower dot in the ellipse). Coupling signals of interesting minor peaks are marked with red rectangles.

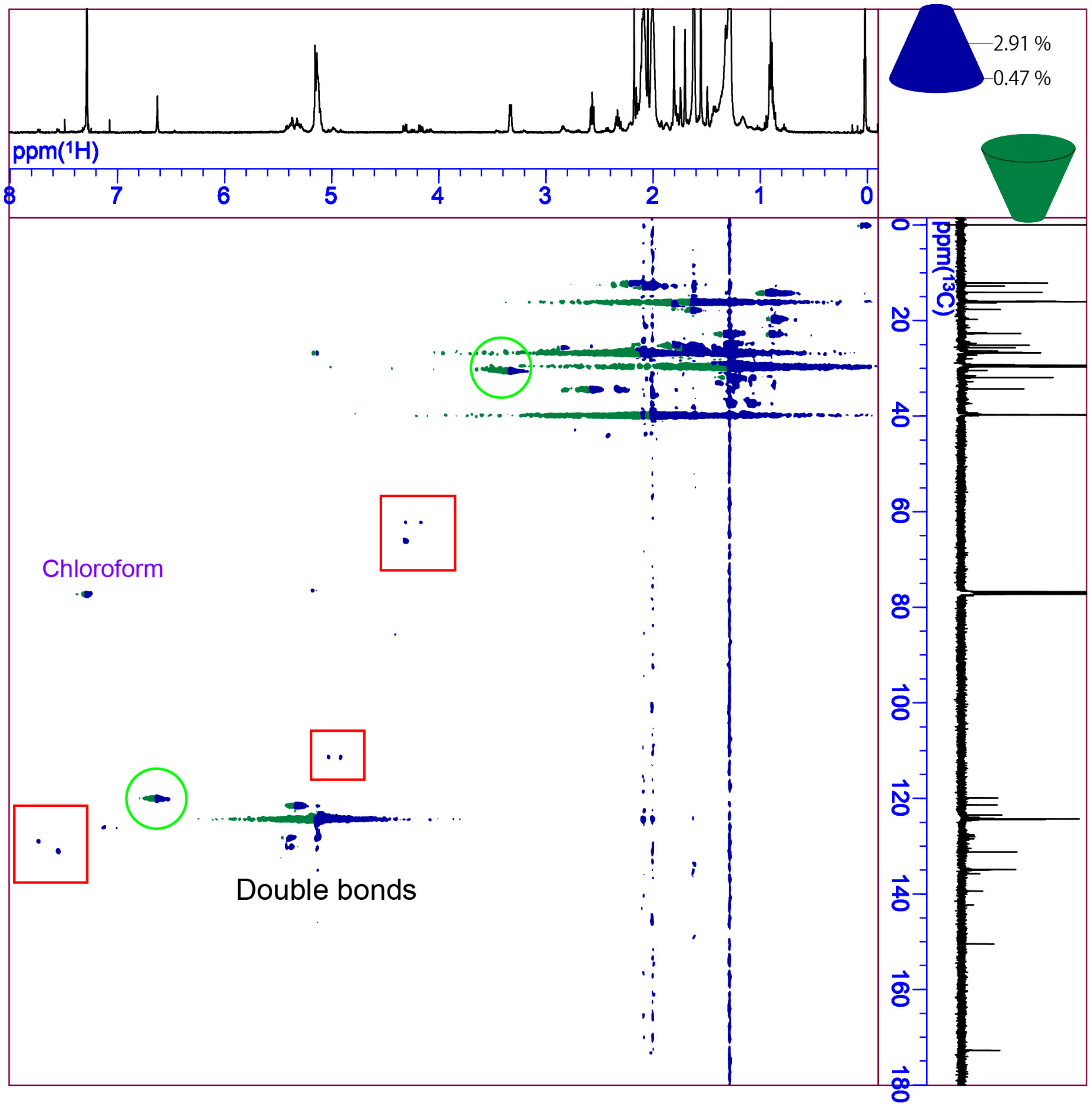


**Supplementary Fig. S2** HSQC analysis of *Synechocystis* TAGL. Coupling of directly linked $^{13}C$ – $^{1}H$ is detected as blue signals. The signals characteristic of the plastoquinonol structure are marked with green circles. They are prenyl 1a (C: 30.4 ppm, H: 3.30 ppm) and ring 3' (C: 119.9 ppm, H: 6.60 ppm). Coupling signals of interesting minor peaks are marked with red rectangles.

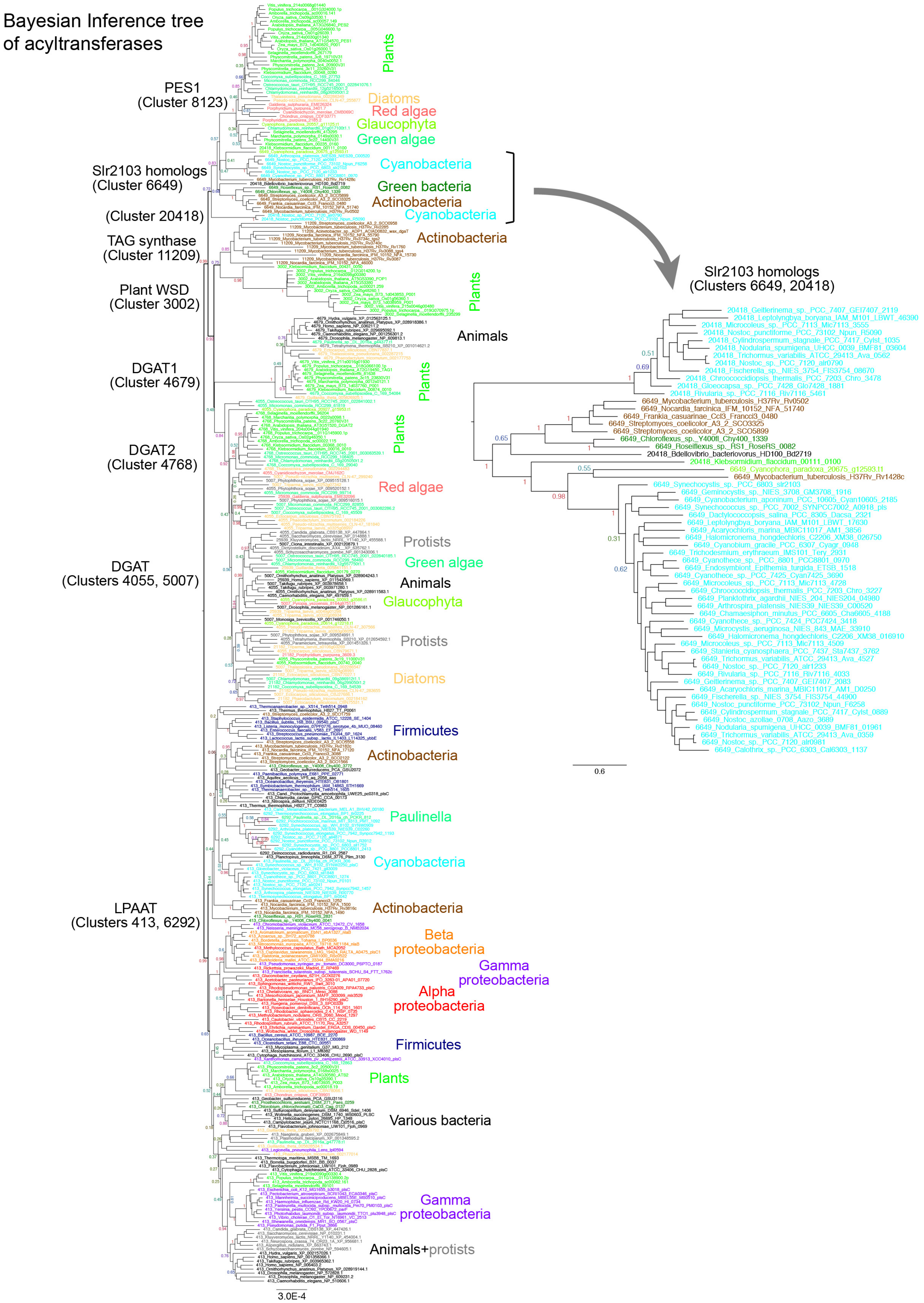
Bayesian Inference tree
of acyltransferases
PES1
(Cluster 8123)
Plants
Diatoms
Red algae
Glaucophyta
Green algae
Slr2103 homologs
(Cluster 6649)
Cyanobacteria
Green bacteria
Actinobacteria
(Cluster 20418)
Cyanobacteria
TAG synthase
(Cluster 11209)
Actinobacteria
Plant WSD
(Cluster 3002)
Plants
Animals
DGAT1
(Cluster 4679)
Plants
DGAT2
(Cluster 4768)
Plants
Red algae
Protists
Green algae
DGAT
(Clusters 4055, 5007)
Animals
Glaucophyta
Protists
Diatoms
Firmicutes
Actinobacteria
Paulinella
Cyanobacteria
LPAAT
(Clusters 413, 6292)
Actinobacteria
Beta
proteobacteria
Gamma
proteobacteria
Alpha
proteobacteria
Firmicutes
Plants
Various bacteria
Gamma
proteobacteria
Animals+protists
3.0E-4
Slr2103 homologs
(Clusters 6649, 20418)
20418_Geitlerinema_sp._PCC_7407_GEI7407_2119
20418_Leptolyngbya_boryana_IAM_M101_LBWT_46390
20418_Microcoleus_sp._PCC_7113_Mic7113_3555
20418_Nostoc_punctiforme_PCC_73102_Npun_R6060
20418_Cylindrospermum_stagnale_PCC_7417_Cylst_1035
20418_Nodularia_spumigena_UHCC_0039_BMF81_03604
20418_Trichormus_variabilis_ATCC_29413_Ava_0562
20418_Nostoc_sp._PCC_7120_alr0790
20418_Fischerella_sp._NIES_3754_FIS3754_08670
20418_Chroococcidiopsis_thermalis_PCC_7203_Chro_3478
20418_Gloeocapsa_sp._PCC_7428_Glo7428_1881
20418_Rivularia_sp._PCC_7116_Riv7116_5461
6649_Mycobacterium_tuberculosis_H37Rv_Rv0502
6649_Nocardia_farcinica_IFM_10152_NFA_51740
6649_Frankia_casuarinae_CcI3_Francci3_0480
6649_Streptomyces_coelicolor_A3_2_SCO3325
6649_Streptomyces_coelicolor_A3_2_SCO5899
6649_Chloroflexus_sp._Y400fl_Chy400_1339
6649_Roseiflexus_sp._RS1_RoseRS_0082
20418_Bdellovibrio_bacteriovorus_HD100_Bd2719
20418_Klebsormidium_flaccidum_00111_0100
6649_Cyanophora_paradoxa_20675_g12593.t1
6649_Mycobacterium_tuberculosis_H37Rv_Rv1428c
6649_Synechocystis_sp._PCC_6803_slr2103
6649_Geminocystis_sp._NIES_3708_GM3708_1916
6649_Cyanobacterium_aponinum_PCC_10605_Cyan10605_2185
6649_Synechococcus_sp._PCC_7002_SYNPCC7002_A0018_pls
6649_Dactylococcopsis_salina_PCC_8305_Dacsa_2321
6649_Leptolyngbya_boryana_IAM_M101_LBWT_17630
6649_Acaryochloris_marina_MBIC11017_AM1_3856
6649_Halomicronema_hongdechloris_C2206_XM38_026750
6649_Cyanothece_sp._PCC_8801_PCC8801_0940
6649_Cyanobium_gracile_PCC_6307_Cyagr_0948
6649_Trichodesmium_erythraeum_IMS101_Tery_2931
6649_Cyanothece_sp._PCC_8801_PCC8801_3600
6649_Endosymbiont_Epithemia_turgida_ETSB_1518
6649_Cyanothece_sp._PCC_7425_Cyan7425_3690
6649_Microcoleus_sp._PCC_7113_Mic7113_4728
6649_Chroococcidiopsis_thermalis_PCC_7203_Chro_3227
6649_Planktothrix_agardhii_NIES_204_NIES204_04980
6649_Arthrospira_platensis_NIES39_NIES39_C00520
6649_Chamaesiphon_minutus_PCC_6605_Cha6605_4188
6649_Cyanothece_sp._PCC_7424_PCC7424_3418
6649_Microcystis_aeruginosa_NIES_843_MAE_33910
6649_Halomicronema_hongdechloris_C2206_XM38_016910
6649_Microcoleus_sp._PCC_7113_Mic7113_4509
6649_Stanieria_cyanosphaera_PCC_7437_Sta7437_3762
6649_Trichormus_variabilis_ATCC_29413_Ava_4527
6649_Nostoc_sp._PCC_7120_alr1233
6649_Rivularia_sp._PCC_7116_Riv7116_4033
6649_Geitlerinema_sp._PCC_7407_GEI7407_2083
6649_Acaryochloris_marina_MBIC11017_AM1_D0250
6649_Fischerella_sp._NIES_3754_FIS3754_44900
6649_Nostoc_punctiforme_PCC_73102_Npun_F6258
6649_Cylindrospermum_stagnale_PCC_7417_Cylst_0889
6649_Nostoc_azollae_0708_Aazo_3689
6649_Nodularia_spumigena_UHCC_0039_BMF81_01961
6649_Trichormus_variabilis_ATCC_29413_Ava_0359
6649_Nostoc_sp._PCC_7120_alr0981
6649_Calothrix_sp._PCC_6303_Cal6303_1137
0.6

**Supplementary Fig. S3** Bayesian Inference (BI) phylogenetic tree of acyltransferases. BI calculation was performed according to Sato and Awai (2017) and Sato (2020). The MrBayes software version 3.2.7a was run in the supercomputer system of the Human Genome Center at the University of Tokyo. The cluster data were obtained from the Gclust database (dataset Gclust2021) at http://gclust.c.u-tokyo.ac.jp/. Major parameters were: aamodelpr=fixed(lg), rates = invgamma, ratepr = variable, ngen = 20,000,000. Major groups of organisms are color-coded. The large tree (left) was constructed with 323 sequences having 190 sites. The small tree (right) consisting of mainly cyanobacterial homologs of Slr2103 was constructed with 59 sequences having 263 sites.
